# Aurora B and C kinases regulate prophase exit and chromosome segregation during spermatogenesis

**DOI:** 10.1101/868836

**Authors:** Stephen R. Wellard, Karen Schindler, Philip Jordan

## Abstract

Precise control of chromosome dynamics during meiosis is critical for fertility. A gametocyte undergoing meiosis coordinates formation of the synaptonemal complex (SC) to promote efficient homologous chromosome recombination. Subsequent disassembly of the SC is required prior to meiotic divisions to ensure accurate segregation of chromosomes. We examined the requirements of the mammalian Aurora kinases (AURKA, B, and C) during SC disassembly and chromosome segregation using a combination of chemical inhibition and gene deletion approaches. We find that both mouse and human spermatocytes fail to disassemble SC lateral elements when AURKB and AURKC are inhibited. Interestingly, both *Aurkb* conditional knockout and *Aurkc* knockout spermatocytes successfully progress through meiosis and mice are fertile. In contrast, *Aurkb*, *Aurkc* double knockout spermatocytes failed to coordinate disassembly of SC lateral elements with chromosome segregation, resulting in delayed meiotic progression, spindle assembly checkpoint failure, chromosome missegregation, and abnormal spermatids. Collectively, our data demonstrates that AURKB and AURKC functionally compensate for one another ensuring successful mammalian spermatogenesis.

**SUMMARY:** Chemical inhibition and gene deletion approaches show that Aurora B and Aurora C have overlapping functions that ensure timely disassembly of lateral element components of the synaptonemal complex in mouse and human spermatocytes and ensure accurate chromosome segregation during meiosis.

## IV. INTRODUCTION

Crossover formation is essential to mediate bidirectional segregation of homologs during meiosis I. To mediate interactions between homologous chromosomes and facilitate completion of homologous recombination, meiotic cells assemble a proteinaceous scaffold termed the synaptonemal complex (SC). The SC not only acts as a bridge between homologs, but it mediates protein interactions and signaling pathways required for meiotic progression (Rog et al., 2017).

The SC is a zipper-like tripartite protein complex comprised of two lateral elements (LEs) and a central region. LE components include meiosis-specific axial proteins (SYCP2 and SYCP3) and cohesin complexes, which collectively form a core between each pair of sister chromatids. The LEs of a pair of homologous chromosomes are bridged together by a series of transverse filaments (SYCP1) and central element proteins (SYCE1-3 and TEX12). Synapsis and crossover recombination between homologous chromosomes is completed by the pachytene sub-stage of meiotic prophase (Cole et al., 2012). Upon completion of HR, spermatocytes are licensed to progress through the prophase to metaphase I (G2/MI) transition (Hochwagen and Amon, 2006). The SC is disassembled in a coordinated manner. Initially, transverse filament and central element proteins are removed during diplonema, allowing homologs to begin disengaging. During diakinesis, LE proteins and cohesins are depleted from the axis but are retained at the kinetochore region (Ishiguro et al., 2011). Final stages of SC disassembly are coordinated with other aspects of the transition to prometaphase/metaphase I, such as chromatin condensation and formation of bivalents (Clemons et al., 2013).

Treatment of mammalian spermatocytes with okadaic acid (OA), a PP1 and PP2 phosphatase inhibitor, stimulates spermatocytes to undergo a number of hallmarks of the G2/MI transition including SC disassembly and formation of condensed bivalents (Wiltshire et al., 1995). Based on these findings, cell-cycle kinases are implicated to play an important role in regulating SC disassembly. In mice, inhibition of cyclin-dependent kinases (CDKs) prevents LE disassembly (Sun et al., 2010). In addition, polo-like kinase 1 (PLK1) directly phosphorylates SYCP1, TEX12 and SYCE1 proteins, and inhibition of these modifications during an OA-induced prophase exit prevents SC disassembly (Jordan et al., 2012).

Aurora kinases (AURKs) are implicated in regulating SC dynamics during meiosis. In budding yeast there is a single AURK, Ipl1, which promotes efficiency of SC disassembly and integrates chromosome restructuring events with cell cycle progression (Jordan et al., 2009; Newnham et al., 2013). Mammalian spermatocytes, however, express three AURK paralogs (AURKA, B, and C) (Tang et al., 2006). Despite a high degree of sequence similarity between the catalytic kinase domains, the three mammalian AURKs display unique functions and localizations. AURKA localizes to centrosomes and spindle microtubules and plays important roles in ensuring bipolar spindle formation (Sugimoto et al., 2002). Both AURKB and AURKC can function as the catalytic subunit of the chromosome passenger complex (CPC), which is composed of the scaffold inner centromere protein (INCENP), and two regulatory subunits survivin, and borealin (Slattery et al., 2008). The CPC localizes to pericentromeric heterochromatin and along chromosome arms beginning at diplonema, and concentrates at kinetochores during diakinesis (Parra et al., 2003). Localization of the CPC in mouse spermatocytes during the G2/MI transition suggests AURKB/C are at the right place at the right time to regulate SC dynamics (Tang et al., 2006). Here, we have used AURKA and AURKB/C inhibitors, mouse mutant models, and human spermatocytes to further investigate the roles of these AURK paralogs in regulating SC and chromosome dynamics.

## V. RESULTS AND DISCUSSION

### Inhibition of AURKB/C results in impaired LE disassembly

Mouse spermatocytes were induced to undergo the G2/MI transition via treatment with OA during a short-term culture. Juvenile mice 18 days post-partum (dpp), undergoing the semi-synchronous first wave of spermatogenesis, were used to obtain an enriched pool of pachytene-stage spermatocytes. To assess LE disassembly, spermatocytes were monitored by immunolabeling SYCP3 on chromatin spreads following 5 hours of culture. Without addition of OA, the G2/MI transition does not occur in cultured spermatocytes. In contrast, cells treated with OA progressed to a prometaphase-like state in which LEs disassembled from the chromosome axis, remaining only at kinetochores (Fig. 1A and B).

**Figure 1.**
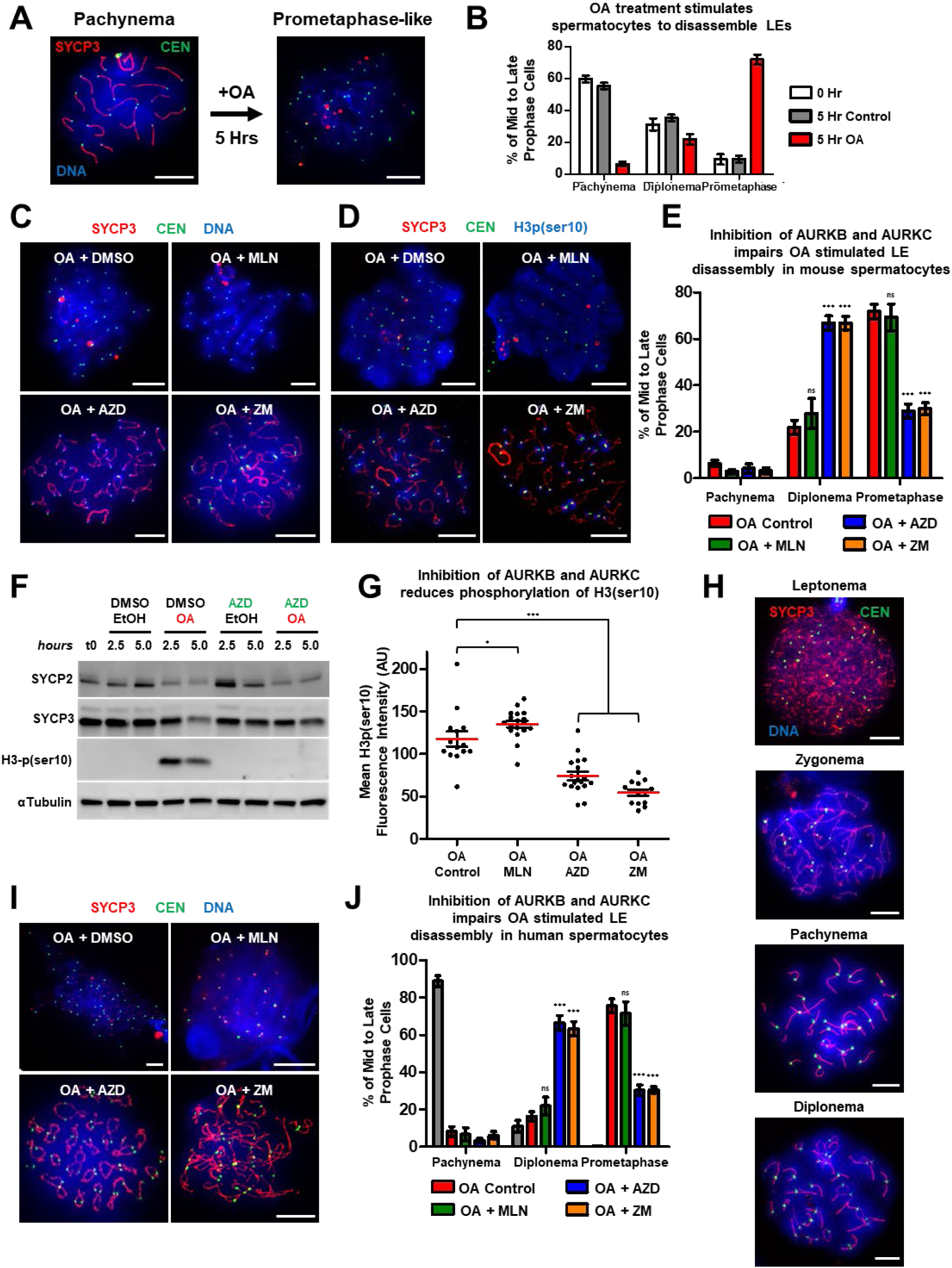
Inhibition of AURKB and AURKC prevents LE disassembly during OA induced G2/MI transition. SC disassembly was monitored by chromatin spread preparations immunolabeled with SYCP3 (red), centromeres (green), and stained with DAPI (blue). **(A)** Pachytene stage chromatin spread prior to chemical treatment and following a 5-hour OA [4µM] treatment. **(B)** Progression through prophase I substages following OA treatment. **(C, D)** Representative images following culture in the presence of OA [4µM] and AURK inhibitors [5µM] immunolabeled with SYCP3 (red), centromeres (green), and stained with DAPI (C) or immunolabeled with H3p(ser10) (blue) (D). **(E)** Progression through prophase I substages following OA and AURK inhibitor treatment. **(F)** Mean nuclear H3p(ser10) signal of treatment conditions shown in (D). **(G)** Western blots of SYCP1-3, SYCE1, H3-p(ser10), and alpha-Tubulin following culture in the presence or absence of OA [4µM] and AZD [5µM]. (**H)** Chromatin spreads from STA-PUT purified human spermatocytes. **(I)** Representative human chromatin spread preparations following a 5-hour OA treatment [4µM] in the presence of AURK inhibitors [5µM]. **(J)** Progression of human spermatocytes following culture with OA, and AURK inhibitors. Error bars in (B, E, G, and J) show mean ± SEM. P values (two-tailed Student’s t-test) comparing each AURK inhibitor treatment to the relevant OA control are indicated by n.s. (not significant), *P<0.05, ***P<0.0001. Scale bar: 10μm. See materials and methods.

To study the role of AURKs on SC disassembly, we used small molecule inhibitors with various affinities for the three AURK paralogs. Specifically, the pan-AURK inhibitor ZM447439 (ZM) (Ditchfield et al., 2003), the AURKA inhibitor MLN8054 (MLN) (Manfredi et al., 2007), and the AURKB/C inhibitor AZD1152 (AZD) (Yang et al., 2007) were used. Treatment of pachytene-stage spermatocytes with AURK inhibitors alone for 5 hours did not induce meiotic progression, SC disassembly, or centromeric cohesion aberrancies (Fig. S1A-C). As previously demonstrated, treatment of spermatocytes with OA in combination with ZM significantly impaired LE disassembly (Sun and Handel, 2008). Interestingly, the same defect in LE disassembly was observed during OA + AZD treatment, while no defect in desynapsis was observed under OA + MLN conditions (Fig. 1C-E). In addition to SYCP3, the meiotic-specific cohesin component REC8 was also retained along the axis under OA + AZD and OA + ZM treatment (Fig. S1D and E). However, sororin, a cohesin protector, which localizes to the SC in a synapsis-dependent manner (Gómez et al., 2016; Jordan et al., 2017), was not retained along the axis under any AURK inhibitor treatment (Fig. S1F and G). We then assessed total levels of LE proteins by western blotting and found that they remained higher after OA + AZD treatment compared to OA treatment (Fig. 1F). These findings suggest that AURKB and AURKC activity regulates LE disassembly, and that AURKA is not required for this event during the G2/MI transition.

Phosphorylation of histone H3 at serine residue 10 (pH3(ser10)) precedes entry into metaphase and chromosome condensation (Hendzel et al., 1997). All three AURKs can catalyze this histone modification (Sugiyama et al., 2002; Li et al., 2004). Analysis of mouse spermatocytes cultured in the presence of OA confirmed that inhibition of AURKB and AURKC with AZD abrogated pH3(ser10) levels from chromosome arms (Fig. 1D, F, and G). pH3(ser10) signal was not diminished within pericentromeric heterochromatin following AURKB and C inhibition, or with pan-AURK inhibition (Fig. 1D). This result suggests that AURKB and AURKC activity is required for pH3(ser10) along chromosome arms, but other kinases may be capable of this modification within pericentromeric heterochromatin regions. Other kinases with documented phospho-H3ser10 activity include CHK1, PKCα, PAK1, and VRK1 (Zhang et al., 2017; Liokatis et al., 2012).

We then extended our analysis of AURK requirements during SC disassembly to human. Human spermatocytes were isolated via STA-PUT density sedimentation and SC dynamics were monitored by SYCP3 immunolabeling of chromatin spread preparations (Fig. 1H). SC assembly and synapsis occurred normally in human spermatocytes obtained from each donor (Fig. 1H, S2A). Treatment of human pachytene-stage enriched spermatocytes with OA induced the G2/MI transition (Fig. 1I and J). Human spermatocytes treated with OA and ZM or AZD, but not MLN, displayed defective LE disassembly, demonstrating that AURKB and AURKC activity is required for LE disassembly in human spermatocytes. These data indicate that a requirement of AURKB/C for exiting meiotic prophase in spermatocytes is conserved between mouse and human.

### Genetic deletion of *Aurkb* or *Aurkc* does not impact the G2/MI transition or the first meiotic division

To further study their roles of AURKB/C during mammalian spermatogenesis, we used germ-cell specific *Aurkb* conditional knockout (cKO) and *Aurkc* knockout (KO) mice (Kimmins et al., 2007; Fernández-Miranda et al., 2011). We tested deleting *Aurkb* using the *Stra8-Cre*, which resulted in severely disrupted entry into spermatogenesis (Fig. S2B-G). Instead, optimal conditional deletion of *Aurkb* was driven by *Spo11-Cre*, which is expressed in spermatocytes shortly after meiotic entry (Lyndaker et al., 2013) (Fig. S3A, and B). Hematoxylin and eosin (H&E) stained testis sections of control, *Aurkb* cKO, and *Aurkc* KO mice showed no histological aberrancies compared to control mice, were fertile, and had similar litter sizes compared to controls when mated to wild type females (Fig. 2A and B). We analyzed SC dynamics during meiotic progression and did not observe any defects in *Aurkb* cKO and *Aurkc* KO mice (Fig. 2C-E, S3C and D). These results demonstrate that the critical axis restructuring events that occur during the G2/MI transition are not affected by the absence of either AURKB or AURKC.

**Figure 2.**
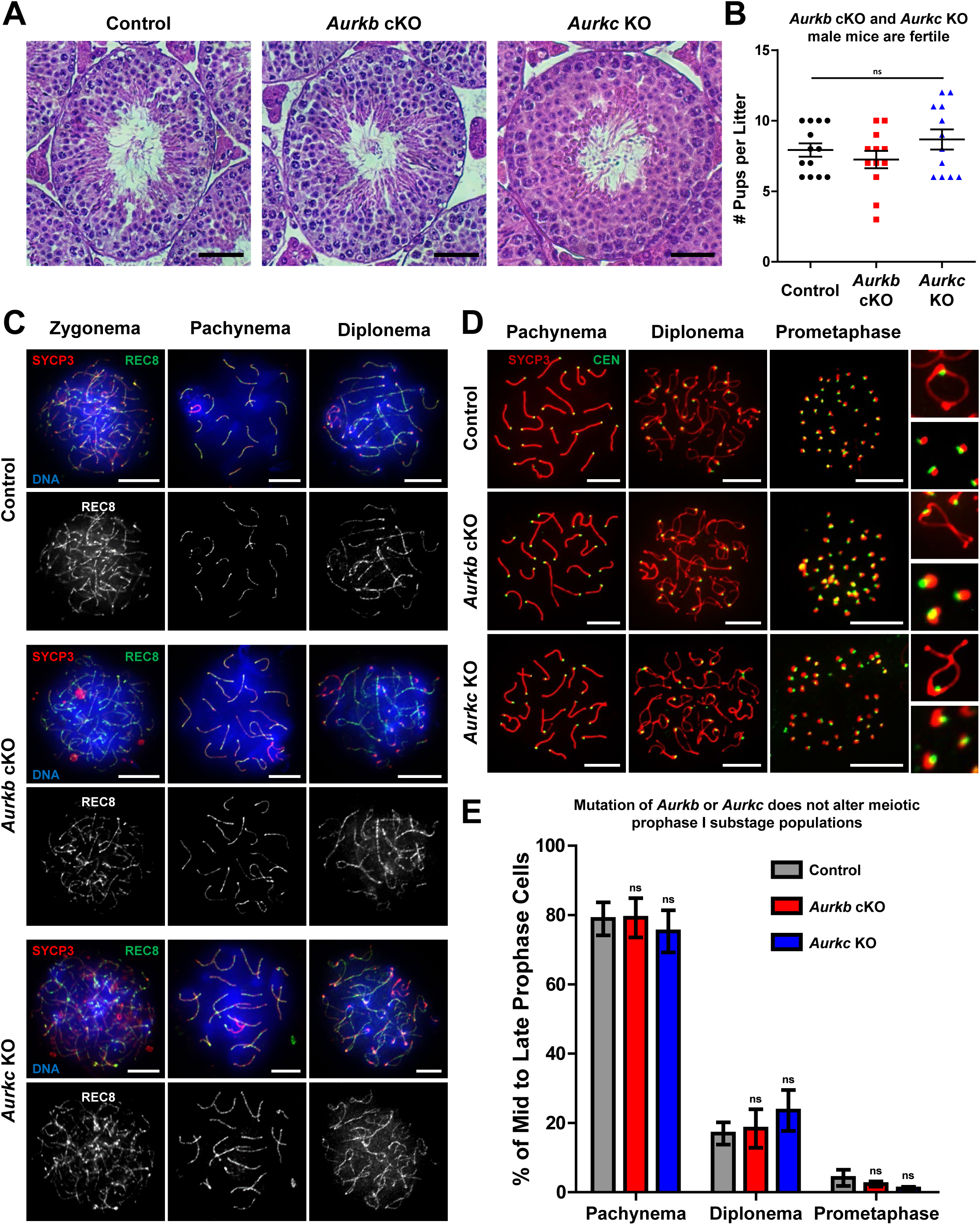
Conditional mutation of *Aurkb* or knockout of *Aurkc* does not result in G2/MI transition aberrancies. **(A)** H&E stained testis sections of adult control, *Aurkb* cKO, and *Aurkc* KO mice. Scale bar: 50µm. **(B)** Fertility tests of male control, *Aurkb* cKO, and *Aurkc* KO mice. **(C, D)** Representative chromatin spread preparations of mid prophase spermatocytes in control, *Aurkb* cKO, and *Aurkc* KO mice stained with DAPI (blue) immunolabeled with SYCP3 (red), REC8 (green) (C) or centromeres (green) (D). **(E)** Prophase I substage distribution in juvenile (18 dpp) control, *Aurkb* cKO, and *Aurkc* KO mice. Scale bar: 10μm. See materials and methods.

We then assessed changes in AURK subcellular localization. Neither AURKB kinetochore localization in *Aurkc* KO, nor AURKC localization *Aurkb* cKO spermatocytes were altered (Fig. 3A). Similarly, the expected localization of AURKA to the centrosome and spindle pole was observed in both mutant mice. This is in contrast to what has been reported for *Aurkc* KO oocytes, where AURKA localizes to spindle poles and chromosomes (Nguyen et al., 2018a). Protection of centromeric sister chromatid cohesion by shugoshin-2 (SGOL2), a known AURKB substrate, during the first meiotic division is critical to ensure sister chromosome mono-orientation and prevent aneuploidy (Llano et al., 2008). In both *Aurkb* cKO and *Aurkc* KO spermatocytes, SGOL2 protein is loaded during the first meiotic division, equivalent to controls (Fig. 3B). During *C. elegans* and *D. melanogaster* meiosis, Aurora kinases are required to remove cohesin from chromosome arms (Rogers et al., 2002; Resnick et al., 2006). However, localization of REC8 was unchanged in *Aurkb* cKO and *Aurkc* KO relative to control mice (Fig. 3C). Quantification of spindle morphology reveals no defects at metaphase I, and spermatocytes from both *Aurkb* and *Aurkc* mutant mice progress to form bipolar spindles (Fig. 3D). These results suggest that deletion of either AURKB or AURKC does not disrupt spermatogenesis and these two kinases functionally compensate for one another within the testis. In oocytes, AURKA and AURKB can compensate for loss of AURKC, but AURKC cannot compensate for loss of AURKB (Nguyen et al., 2018a). These differences highlight sexual dimorphism in Aurora kinase regulation.

**Figure 3.**
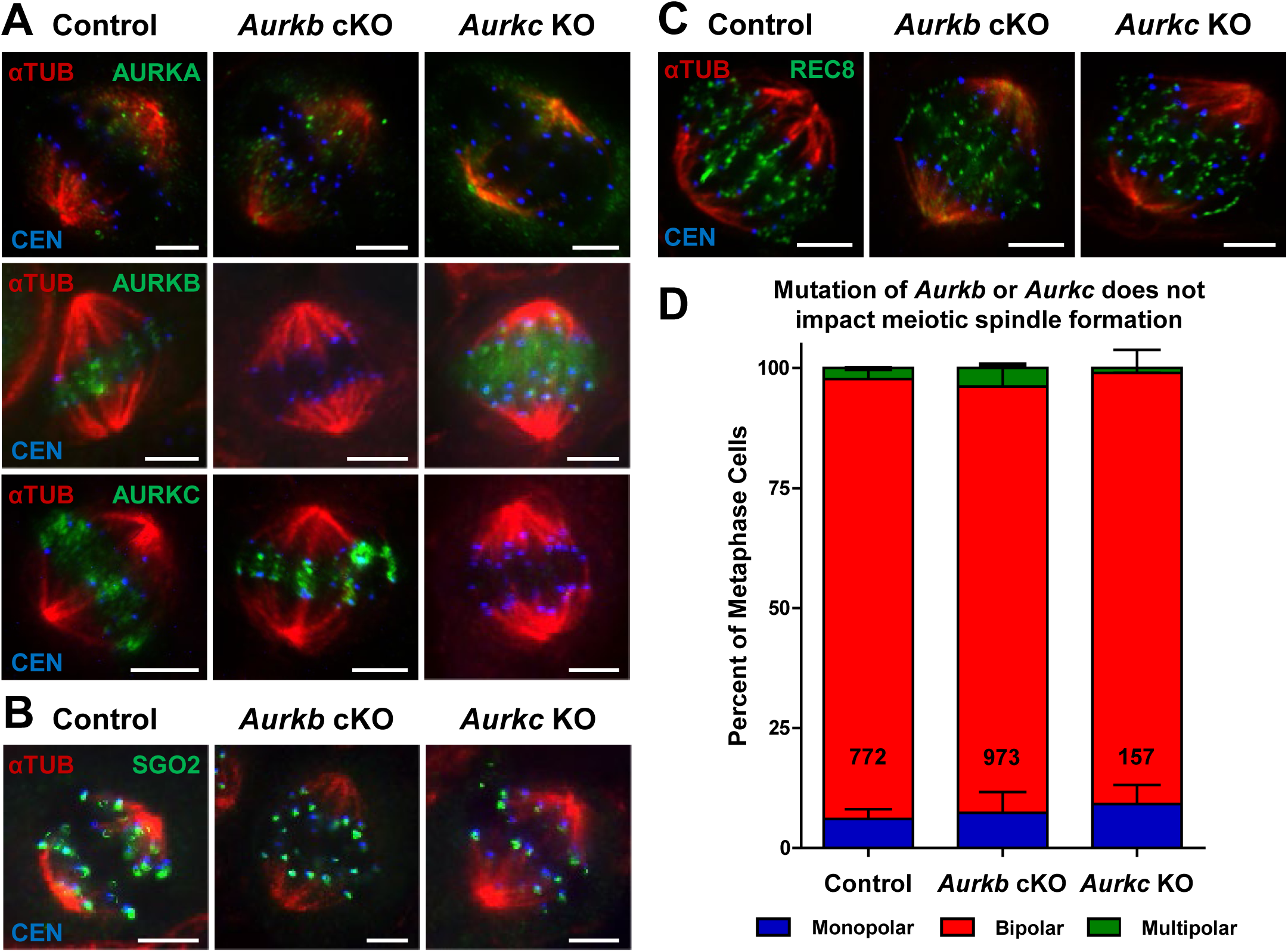
Deletion of AURKB or AURKC does not impact the first meiotic division. **(A)** Spermatocytes from tubule squash preparations were immunolabeled with antibodies against alpha-tubulin (red), centromeres (blue), and the three mammalian AURKs (green), SGO2 **(B)**, and REC8 **(C)**. Scale bar: 5μm. **(D)** Quantification of spindle polarity during the first meiotic division for control, *Aurkb* cKO, and *Aurkc* KO mice. Mean of three biological replicates and total number of cells assessed is indicated. Error bars in (D) represent mean ± SEM. P values (two-tailed Student’s t-test) were not significant. See materials and methods.

Expression of a kinase inactive AURKB mutant led to impaired spermatogenesis with multinucleated spermatocytes (Kimmins et al., 2007). Similarly, mutation in human *AURKC* that generates a kinase inactive truncation results in polyploid sperm (Dieterich et al., 2007). Thus, mutation of the kinase domain in AURKB and AURKC had a greater impact on spermatogenesis than the cKO and KO approaches used here. Both AURKB and AURKC can bind to INCENP and function as the catalytic subunits of the CPC; however, this binding cannot occur simultaneously (Sasai et al., 2016). Mutant forms of either AURKB or AURKC may deplete the pool of active CPC available to a developing spermatocyte. Therefore, mutations that affect the catalytic function of AURKB or AURKC could display a dominant negative effect, as has been observed in oocytes (Balboula and Schindler 2014; Fellmeth et al. 2016; Nguyen et al. 2017), and interfere with CPC function.

### Deletion of both AURKB and AURKC results in LE disassembly defect

Because AURKB and AURKC appear to have redundant functions in spermatogenesis, we assessed spermatocytes from mice lacking both kinases. Histological analyses of testis sections obtained from adult *Aurkb/c* double knockout (dKO) mice showed severe disruption of meiotic progression, with accumulation of primary spermatocytes (Fig. 4A-D). Immunolabeled cryosections of seminiferous tubules from mice completing the first wave of spermatogenesis revealed an accumulation of prophase spermatocytes in *Aurkb/c* dKO mice (Fig. 4B, C, and S3E). Cells with linear stretches of SYCP3 and condensed γH2AX-rich XY chromosome pairs accumulated in *Aurkb/c* dKO mutant tubules (Fig. 4B and D). Assessment of chromatin spread preparations demonstrated that *Aurkb/c* dKO spermatocytes retain linear stretches of SYCP3 on fully condensed bivalent chromosomes, which contrasts with control spermatocytes where SYCP3 is only present at kinetochores (Fig. 4E). These results demonstrate that LE disassembly is perturbed in *Aurkb/c* dKO spermatocytes.

**Figure 4.**
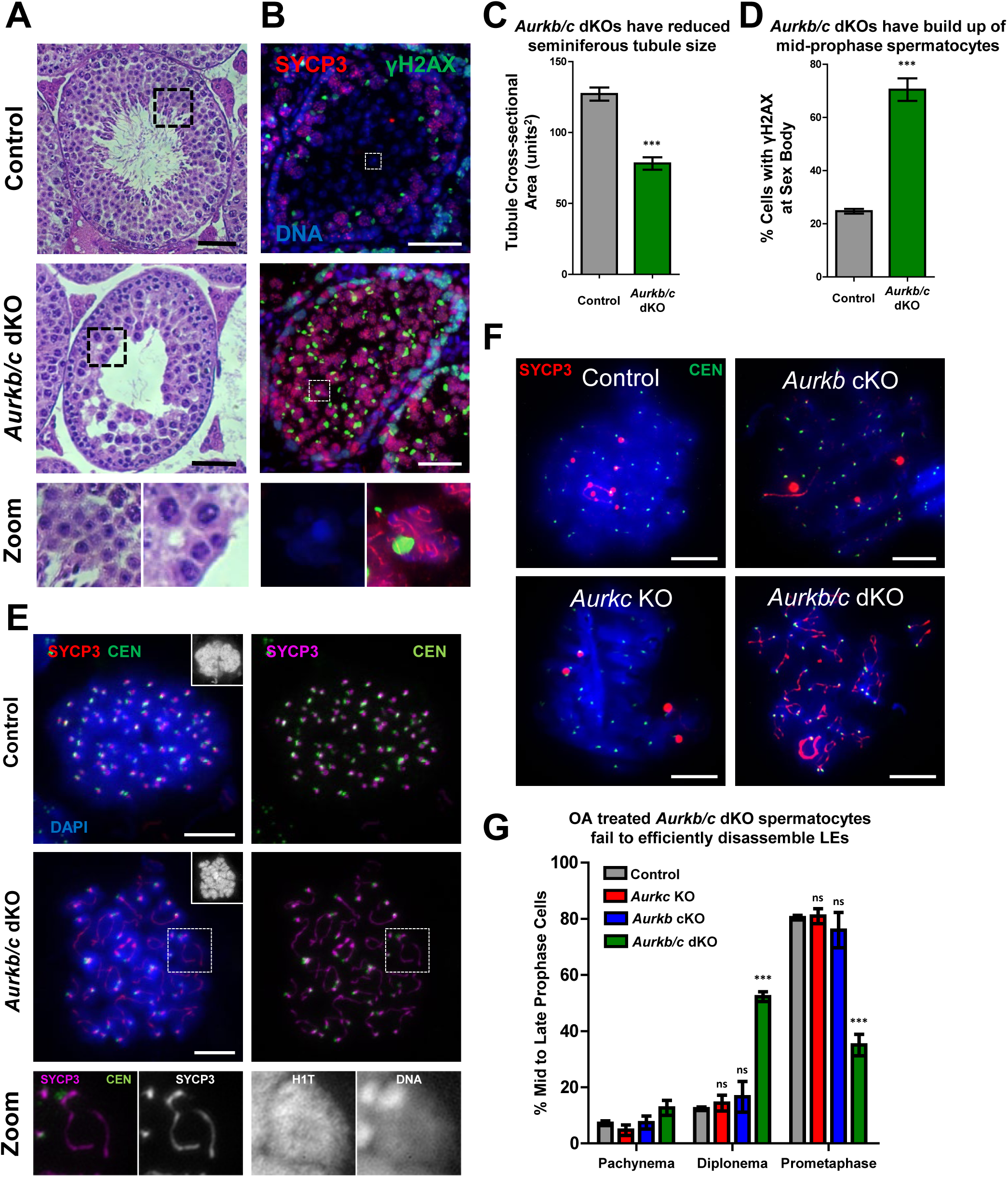
Deletion of both AURKB and AURKC results in an LE disassembly defect. **(A)** H&E stained testis sections of adult control, and *Aurkb/c* dKO mice. Scale bar 50µm. **(B)** Immunohistochemistry of testis cryosections from 33 dpp control and *Aurkb/c* dKO mice immunolabeled with SYCP3 (red), γH2AX (green) and stained with DAPI (blue). Scale bar: 50µm. **(C, D)** Quantification of the tubule area in 33 dpp control and *Aurkb/c* dKO (C), and percent of cells with γH2AX staining at the sex body within seminiferous tubules sections (D). **(E)** Chromatin spread preparations from 18dpp control and *Aurkb/c* dKO mice immunolabeled with SYCP3 (red), centromeres (green), and stained with DAPI. **(F)** Representative chromatin spread preparations of control, *Aurkb* cKO, *Aurkc* KO, and *Aurkb/c* dKO mice after a 5-hour treatment with OA [4µM]. Scale bar: 10µm. **(G)** Quantification of cells with linear SYCP3 stretches following OA treatment. Error bars in (C), (D), and (G) show mean ± SEM. P values (two-tailed Student’s t-test) comparing control and mutant mice indicated by n.s. (not significant), ***P<0.0001. See materials and methods.

Spermatocytes isolated from *Aurkb/c* dKO mice were treated with OA to determine whether completion of the G2/MI transition could be chemically induced outside the context of the seminiferous tubule. *Aurkb/c* dKO spermatocytes fail to efficiently disassemble LEs post OA treatment (Fig. 4F and G). In contrast, *Aurkb* cKO and *Aurkc* KO spermatocytes did not display this defect and progressed to prometaphase post OA treatment in a similar manner to control spermatocytes. The enzymatic activity of AURKB/C kinases is therefore required for the efficient removal of LE stretches and the timely completion of desynapsis. Phosphorylation of SC components during the G2/MI transition has previously been reported (Jordan et al., 2012; Fukuda et al., 2012). Interestingly, both SYCP2 and SYCP3 contain putative AURK phosphorylation motifs, and phosphopeptides containing these motifs have been identified in large scale phosphoproteomic studies from mouse testes (Huttlin et al., 2010). Because AURKB and AURKC both localize to the chromosome arms during the G2/MI transition, they are present at the right place at the right time to modify SC components. In future work, identification of AURKB/C targets during spermatogenesis will improve our understanding of the meiotic functions Aurora kinases play during gametogenesis.

### Deletion of both AURKB and AURKC results in chromosome missegregation

Despite abnormal LE disassembly, *Aurkb/c* dKO spermatocytes underwent chromosome segregation. However, the majority of *Aurkb/c* dKO spermatocytes harbored misaligned chromosomes at metaphase I (Fig. 5A-F). The misalignment is not due to mislocalization of AURKA (Fig. 5A), kinetochore proteins crucial for sister chromatid mono-orientation during meiosis I, SGOL2 and MEIKIN (Fig. 5B and C), (Llano et al., 2008; Kim et al., 2015), or premature loss of REC8 cohesins (Fig. 5D). We assessed a spindle assembly checkpoint (SAC) protein, MAD2, which normally localizes to kinetochores during prometaphase, and remains there until ubiquitous bipolar microtubule-kinetochore attachment satisfies the SAC (Lara-Gonzalez et al., 2012). Strikingly, we observe the complete absence of MAD2 from kinetochores in *Aurkb/c* dKO spermatocytes (Fig. 5E). Inhibition studies using ZM have shown that Aurora kinase function is important for MAD2 localization to the kinetochore in mitotic cells (Ditchfield et al., 2003). These results demonstrate that Aurora B and/or C function is required for MAD2 localization to the kinetochore, and, thus, functional SAC during spermatogenesis. We observed meiosis I and meiosis II segregation defects in *Aurkb/c* dKO spermatocytes, many still harboring extensive SYCP3 signal (Fig. 5G-K). Collectively, these defects resulted in the formation of abnormal round spermatids with residual SYCP3 signal.

**Figure 5.**
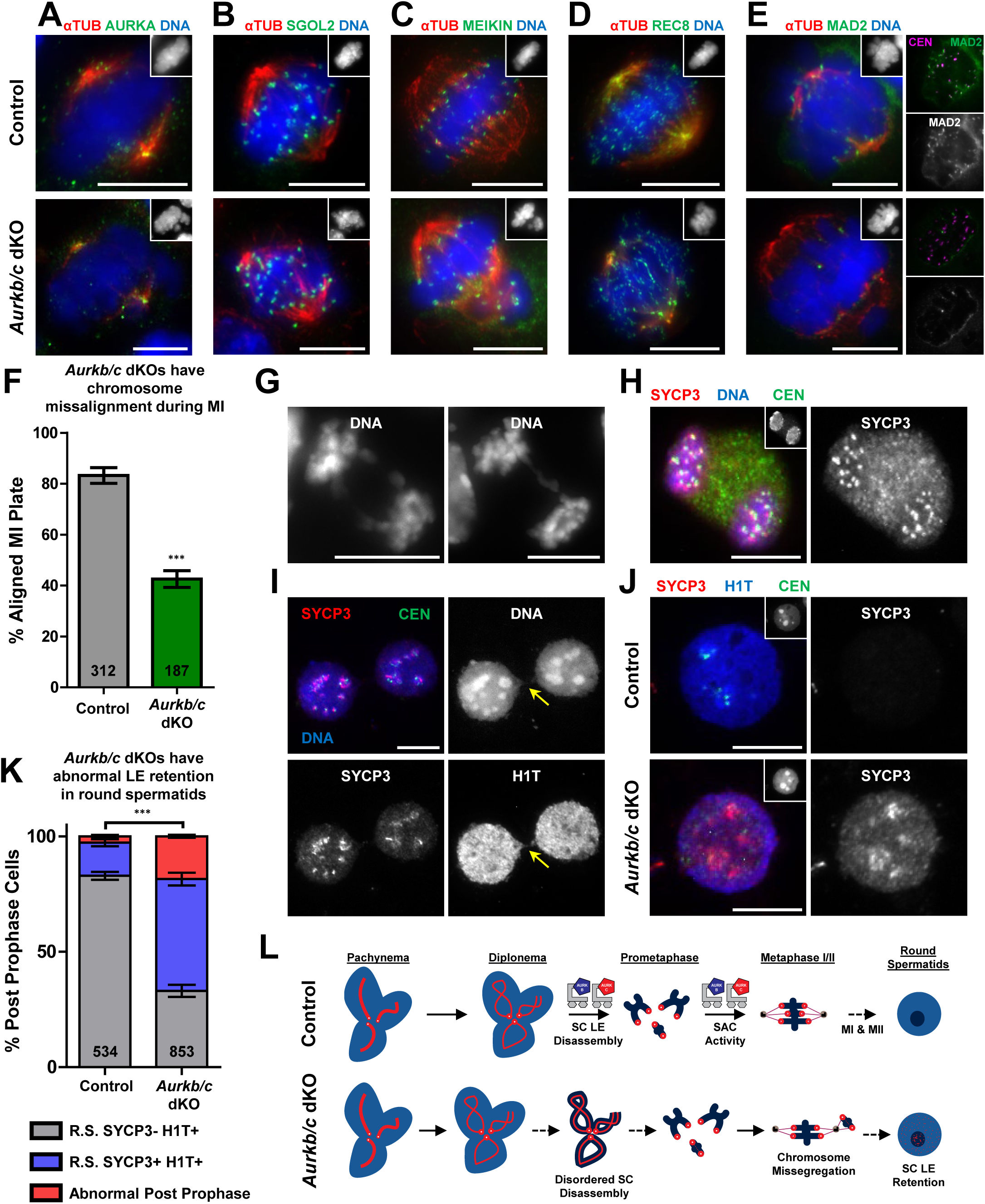
Deletion of both AURKB and AURKC results in SAC failure, chromosome mis-segregation, and formation of round spermatids with residual LE protein. Spermatocytes from tubule squash preparations were stained with DAPI and immunolabeled with antibodies against alpha-tubulin (red), and additional markers (green) including AURKA **(A)**, SGOL2 **(B)**, MEIKIN **(C)**, REC8 **(D)**, and MAD2 **(E)**. **(F)** Quantification of metaphase I plate alignment in control and *Aurkb/c* dKO mice at 23dpp. Mean of three biological replicates and total number of cells assessed is indicated. **(G-J)** Tubule squash (G) and chromatin spread preparations (H-J) from 23dpp control and *Aurkb/c* dKO mice were immunolabeled with SYCP3, centromeres, H1T, and stained with DAPI. DAPI staining is presented as insets in (A-E) and (J), and H1T staining is shown as insets in (H). **(K)** Quantification of post-prophase cell populations in control and *Aurkb/c* dKO mice at 23dpp. Mean of two biological and two technical replicates with the total number of cells assessed indicated. **(L)** Model for the role of AURKB and AURKC during the G2/MI transition in mammalian spermatocytes. Without AURKB and AURKC activity, the coordination and efficiency of LE disassembly is reduced and ultimately decouples LE disassembly with chromatin condensation and cell cycle progression. AURKB and AURKC are required to regulate the spindle assembly checkpoint during both meiosis I and II to ensure accurate chromosome segregation. See materials and methods.

In sum, our data demonstrate that chromosome restructuring events at the axis, and more generally throughout the chromatin, must be coordinated with cell-cycle progression to restrict spermatocytes from entering metaphase I prior to SC disassembly. The budding yeast AURK, Ipl1, blocks cell cycle progression during early meiotic prophase by suppressing S-CDK activity (Newnham et al., 2013). In the context of depleted *Ipl1*, spindle pole body maturation and separation, as well as cell cycle progression are decoupled and cells erroneously progress to metaphase with linear stretches of SC components still present at the axis (Jordan et al., 2009; Newnham et al., 2013). However, mammals express three AURK proteins, where AURKA appears to predominantly function in the cytoplasm during centrosome maturation, and AURKB/C are required for the chromosomal events which take place during the G2/MI transition in males. In contrast, AURKA supports meiosis in the absence of AURKB/C in mouse oocytes by localizing to the kinetochores to become a component of the CPC (Nguyen et al., 2018). Furthermore, deletion of *Aurkb* or *Aurkc* in oocytes results in abnormal meiosis (Schindler et al., 2012; Nguyen et al., 2018). These sexual dimorphisms may be attributed to the spatiotemporal differences between spermatogenesis and oogenesis. Future work must be directed toward determining Aurora kinase substrates at each stage of gametogenesis. For spermatogenesis, this will require *in vivo* synchronization using retinoic acid inhibitor, WIN 18,446 (Hogarth et al., 2013).

---

**Figure S1.**

**Inhibition of AURKs alone does not alter meiotic progression or SC disassembly.**

**(A)** Representative chromatin spread preparations from juvenile (18dpp) C57BL/6J mice prior to treatment and following a 5-hour treatment with AURK inhibitors [5µM] throughout the meiotic prophase I substages of zygonema, pachynema, and diplonema. SC dynamics were monitored by immunolabeling with SYCP3 (red), SYCP1 (green), and centromeres (blue). **(B)** Quantification of the meiotic mid-to-late prophase I substage populations following treatment with AURK inhibitors. Error bars show mean ± SEM. P values (unpaired two-tailed Student’s t-test) comparing control and mutant mice indicated by n.s. (not significant) if above a cutoff of P<.01. **(C)** Maintenance of centromeric pairing in late diplonema was quantified from chromatin spread preparations in control, OA treatment, and AURK inhibitor treatment conditions. Error bars represent mean ± SEM. P values (unpaired two-tailed Student’s t-test) comparing the mean percent paired centromere from control and inhibitor biological replicates indicated by n.s. (not significant), or ***P<0.0001. To further assess SC structure during OA induced disassembly spermatocytes were immunolabeled with SYCP3 (red), the axis component REC8 (green, **D** and **E**), the central element component SORORIN (green, **F** and **G**), and stained with DAPI. Spermatocytes treated with EtOH and DMSO for 5 hours in culture were immunolabeled as described above for (D) and (F), and spermatocytes treated with OA and AURK inhibitors for (E) and (G). Disassembly of SORORIN from the SC proceeded under all AURK inhibitor conditions. Scale bar: 10μm.

**Figure S2.**

**SC synapsis in human spermatocytes and genetic depletion of AURKB using the meiotic specific *Stra8-cre* transgene.**

**(A)** SC assembly and synapsis in human spermatocytes was assessed by immunolabeling with SYCP3 (red), SYCP1 (green), and centromeres (blue). The inset box depicts DAPI staining (white). Scale bars: 10μm. **(B)** Representative image of adult control and *Aurkb Stra8-Cre tg/0* mouse testes. Scale bar: 50mm. **(C)** Relative testis weights were assessed at 10, 14, 20, and >56 (adult) dpp for control (C) and *Aurkb*^*flox/del*^, *Stra8-Cre tg/0* mice (E). Early and persistent defects in relative testis weight suggest severely disrupted spermatogenesis. Hematoxylin and eosin staining of 5-micron thick testis sections of control, and *Aurkb Stra8-Cre tg/0* mice aged 14dpp **(D)** and >56dpp **(E)**. **(F)** Cryo preserved 18dpp testes from control and *Aurkb* cKO (Stra8-Cre) were sectioned and immunolabeled with DAZL and stained with DAPI. **(G)** The ratio of meiotic to post-meiotic cells per tubule cross-section was quantified using DAZL and DAPI staining. Severely depleted seminiferous tubules can be observed as early as 14dpp in *Aurkb*^*flox/del*^, *Stra8-Cre tg/0* mice, indicating germ cell loss prior to meiotic entry. Due to the inefficient Cre excision, patches of germ cells progressing through spermatogenesis can also be observed. Utilization of *Stra8-Cre* for the depletion of AURKB was therefore deemed disadvantageous for the assessment of meiotic SC dynamics. Scale bars: 100μm

**Figure S3.**

**Additional Assessment of *Aurkb* cKO, *Aurkc* KO, and *Aurkb/c* dKO mice.**

**(A)** Protein extracts from STAPUT-isolated pachytene spermatocytes were collected from control and *Aurkb* cKO mice. Isolation via STAPUT density sedimentation resulted in 90% pure mid-prophase spermatocyte enrichment. Western blot analysis was performed for AURKC, AURKB, and alpha-Tubulin as a loading control. **(B)** *Aurkb* cKO male mice were mated to wild-type females and their resulting progeny was assessed for the retention of a *Flox* allele, thereby assessing the efficiency of Cre mediated excision of the floxed allele. Representative chromatin spread preparations of mid prophase spermatocytes in control, *Aurkb* cKO, and *Aurkc* KO mice immunolabeled with SYCP3 (red), HORMAD1 (**C**, green), SORORIN (**D**, green) and stained with DAPI (blue). Scale bar: 10µm. The localization of HORMAD1 and SORORIN was unchanged in spermatocytes isolated from *Aurkb* cKO and *Aurkc* KO mice. **(E)** Depletion of AURKB can be observed via immunohistochemistry of 10μm testis cryosections from 33 dpp control and *Aurkb/c* dKO mice immunolabeled with SYCP3 (red), AURKB (green), and stained with DAPI (blue). Scale bar: 50µm.

## VII. MATERIALS AND METHODS

### a. Mouse and Human Ethics Statement

Mice were bred by the investigators at Rutgers University (Piscataway, NJ) and Johns Hopkins University (Baltimore, MD) under standard conditions in accordance with the National Institutes of Health and U.S. Department of Agriculture criteria and protocols for their care and use were approved by the Institutional Animal Care and Use Committees (IACUC) of Rutgers University and Johns Hopkins University.

Studies involving deidentified donor testes tissues have been reviewed and designated by Johns Hopkins University Bloomberg School of Public Health IRB as “not human subjects research” (IRB No: 00006700).

### b. Mice

To deplete AURKB levels in developing spermatocytes, mice harboring a conditional knockout allele of *Aurkb* (STOCK-*Aurkb*^*tm2.1Mama*^) were used, and have been described previously (Fernández-Miranda et al., 2011). Conditional mutation was achieved by the addition of a hemizygous Cre recombinase transgene under the control of meiosis specific promoters. In this study, both the promoter for *Stra8* (B6.FVB-Tg(*Stra8-iCre)1Reb/LguJ*), and the promoter for *Spo11* (Tg(Spo11-cre)1Rsw) were assessed. *Aurkc* KO mice (B6;129S5-*Aurkc*^*tm1Lex*^) were generated by Lexicon Pharmaceuticals and described previously (Kimmins et al., 2007).

### c. PCR genotyping

Primers used during this study are described in Supplementary Table S1. PCR conditions: 90°C for 2 min; 30 cycles of 90°C for 20 s; 58°C for 30 s; and 72°C for 1 min.

For genotyping of *Aurkc* KO mice, the copy number of *Neo* was quantified by real-time PCR per the manufacturer’s protocol. Briefly, tails were digested in 400 μL of lysis buffer (125 mM NaCl, 40 mM Tris, pH 7.5, 50 mM EDTA, pH 8, 1% (vol/vol) sarkosyl, 5 mM DTT, and 50 μM spermidine) with 6 μL of Proteinase K (Sigma #P4850) for 2 h at 65 °C. After dilution of 1:30 in water, the lysates were boiled for 5 min to denature Proteinase K. Two microliters of the diluted DNA were added to each reaction. Primers to detect *Neo* (F: 5′ CTCCTGCCGAGAAAGTATCCA-3′; R: GGTCGAATGGGCAGGTAG-3′) were used at a final concentration of 300 nM and primers to detect *Csk* (for sample normalization) (F 5′-CTGGC-CATCCGGTACAGAAT-3′; R 5′-TGCAGAAGGGAAGGTCTTGCT-3′) were used at a final concentration of 100 nM. The TAMRA-quenched *Neo* probe (ABI) was conjugated to 6-fluorescein amidite and used at a final concentration of 100 nM and the TAMRA-quenched *Csk* probe was conjugated to VIC and used at a final concentration of 100 nM. The comparative Ct method was used to calculate the *Neo* copy number.

### d. Human testes

Deidentified human organ donor derived human testes utilized in this study were obtained from three donors 43, 23, and 18 years old respectively.

### e. Mouse and human spermatocyte isolation and culturing conditions

Mixed mouse germ cell populations were isolated as described previously (Bellve, 1993; La Salle et al., 2009). Mid-prophase enriched spermatocytes were isolated from 18 dpp mice, undergoing the semi-synchronous first wave of spermatogenesis.

Human mixed germ cell populations were liberated from testis material following two enzymatic digestions, as described previously (Yao et al., 2017; Liu et al., 2015). During the first digestion, seminiferous tubules were isolated by incubation with 2 mg/ml collagenase, and 1 μg/μl DNase I for 15 minutes in an oscillating (100rpm) water bath at 34°C. To release germ cells, the seminiferous tubules were then treated with 3mg/ml collagenase, 2.5 mg/ml hyaluronidase, 2mg/ml trypsin, and 1 μg/μl DNase I for 13 minutes in an oscillating (100rpm) water bath at 34°C. Following the last enzymatic treatment seminiferous tubules were mechanically disrupted using a transfer pipette for 3 minutes on ice. The germ cell mixture was then centrifuged for 7 minutes at 500*g*, resuspended in 25 ml of 0.5% BSA in KRB, and filtered through a 70 μm nitex mesh to create a single cell suspension.

Enriched primary spermatocytes were isolated as using STA-PUT gravity sedimentation as previously described with minor adjustments (La Salle et al., 2009). A density gradient was created by flowing 550 ml of 4% BSA in KRB and 550 ml of 2% BSA in KRB into the 25 ml of cell suspension in 0.5% BSA in KRB. Cells were sedimented for 3 hours prior to elution and fractionation into 12 × 75 mm glass culture tubes. Aliquots from each fraction were assessed to determine the purity of isolated primary spermatocytes, as identified from cell shape and size. Fractions containing abundant (80% pure) primary spermatocytes were pooled, counted, and centrifuged at 500*g* to resuspend at a cell concentration of 2.5×10^6^ cells/ml.

Both mouse and human spermatocytes were cultured at 32°C in 5% CO2 in HEPES (25 mM)-buffered MEMα culture medium (Sigma) supplemented with 25 mM NaHCO3, 5% fetal bovine serum (Atlanta Biologicals), 10 mM sodium lactate, 59 μg/ml penicillin, and 100 μg/ml streptomycin. Spermatocytes were stimulated to undergo the G2/MI transition by a 4 μM okadaic acid (OA) (Sigma) treatment for 5 hours. To assess the role of Aurora kinases on SC disassembly during an OA induced G2/MI transition, spermatocytes were treated with the small molecule inhibitors MLN8054 (Selleck Chemicals), AZD1152 (Sigma), and ZM447439 (Selleck Chemicals) at a concentration of 5 μM.

### f. Mouse and human chromosome spreads

Mouse and human chromatin spread preparations were performed as previously described (Jordan et al., 2012; de Vries et al., 2012). Supplementary Table 2 describes the primary antibodies and their dilutions used in this study. Secondary antibodies conjugated to Alexa 488, 568, or 633 against human, rabbit, and mouse IgG (Life Technologies) were used at 1:500 dilution. Chromatin spreads, and tubule squash preparations were mounted in Vectashield + DAPI (4’, 6-diamidino-2-phenylindole) medium (Vector Laboratories).

To quantify prophase I substage distributions in Figures 1, 2, and 4, chromatin spreads were performed from at least three biological replicates. In addition, the chromatin spreads were performed in duplicate. During analysis the mean percentage of each substage was determined after counting 100-200 cells per technical replicate.

### g. Histology and cryo-sectioning

For histological assessment, mouse testis tissue was fixed in bouins fixative (Ricca Chemical Company) prior to paraffin embedding. Serial sections 5 microns thick were mounted onto slides and stained with hematoxylin and eosin. For cryo-sectioning testis tissue was embedded in O.C.T. compound (Fisher) and frozen on dry ice. Serial sections 5 microns thick were mounted onto slides and immunolabeled with primary and secondary antibodies as described above.

### h. Tubule squash preparations

Mouse tubule squash preparations were performed as previously described (Wellard et al., 2018). Full Z-stack captured images were utilized to manually identify spindle morphology and chromosome alignment.

### i. Western Blot analyses

Protein was extracted from germ cells using RIPA buffer (Santa Cruz) containing 1x protease inhibitor cocktail (Roche). Protein concentration was calculated using a BCA protein assay kit (Pierce), and 20 μg of protein extract was loaded per lane of a 7.5%, 12% or 4-15% gradient SDS PAGE gels (Bio-Rad). To detect proteins >100kDa a 7.5% gel was used, and proteins <100kDa were run on a 12% gel. Protein isolated from STA-PUT enriched pachytene spermatocytes was loaded onto the 4-15% gradient gel. Following protein separation, proteins were transferred to PVDF membranes using Trans-Blot Turbo Transfer System (Bio-Rad). Primary antibodies and dilution used are presented in Supplementary Table 2. For detection of primary antibodies, goat anti-mouse and goat anti-rabbit horseradish peroxidase-conjugated antibodies (Invitrogen) were used as secondary antibodies. Antibody signal was detected via treatment with Bio-Rad ECL western blotting substrate and captured using a Syngene XR5 system.

### j. Microscope image acquisition

Nuclear spread and tubule squash images were captured using a Zeiss CellObserver Z1 linked to an ORCA-Flash 4.0 CMOS camera (Hamamatsu), and histology images were captured using a Zeiss AxioImager A2 with an AxioCam ERc 5s (Zeiss) camera. Images were analyzed with the Zeiss ZEN 2012 blue edition image software and Photoshop (Adobe) was used to prepare figure images.

### k. Statistical Analysis

Student’s t-tests, as indicated in figure legends, were used to evaluate the differences between groups using GraphPad Prism software.

## VIII. ACKNOWLEDGMENTS

The authors thank the Washington Regional Transplant Community for their assistance in obtaining deidentified human testis donations for research, and Marcos Malumbres (CNIO) for providing *Aurkb*^*tm1c*^ mice. Additionally, the authors thank the following researchers for generously providing antibodies for use in this study; Tang K. Tang (Aurora C), José Luis Barbero (SGOL2), Yoshi Watanabe (MEIKIN), Mary Ann Handel (H1T), and Susannah Rankin (Sororin). We would also like to thank Edward Culbertson, Anita Ramachandran, Tianlu Ma, Christopher Shults, Alexandra Nguyen, Amanda Gentilello, and Suzanne Quartuccio for their assistance with mouse genotyping and mouse characterization.

This work was funded by NIGMS grants to P.W.J. (R01GM11755) and K.S. (R01GM112801), Fulbright Distinguished Scholar Award to P.W.J., and training grant fellowship from the National Cancer Institute (NCI, NIH) (CA009110) to S.R.W.

The authors declare no competing financial interests.

## Author contributions

P. Jordan conducted initial experiments, maintained mouse lines, and conceived the project. S. Wellard performed and analyzed all experiments with assistance from P. Jordan. S. Wellard and P. Jordan designed experiments and wrote the manuscript. K. Schindler bred and provided *Aurkc* KO mice, as well as critically reviewed and edited the manuscript.

