## Supplementary figures and images for "Aurora B and C kinases regulate prophase exit and chromosome segregation during spermatogenesis"

### Supplemental Figure S1

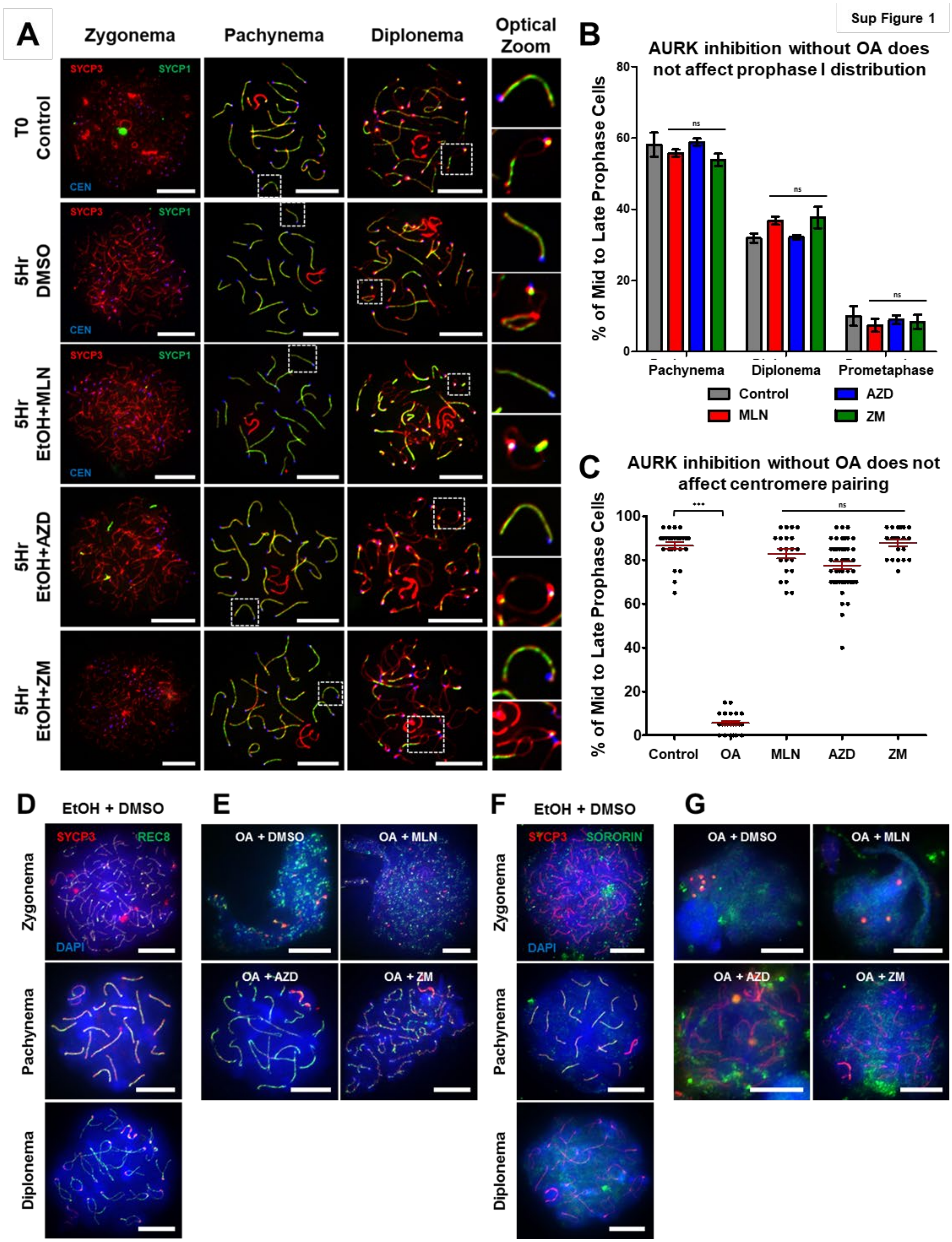

### Supplemental Figure S2

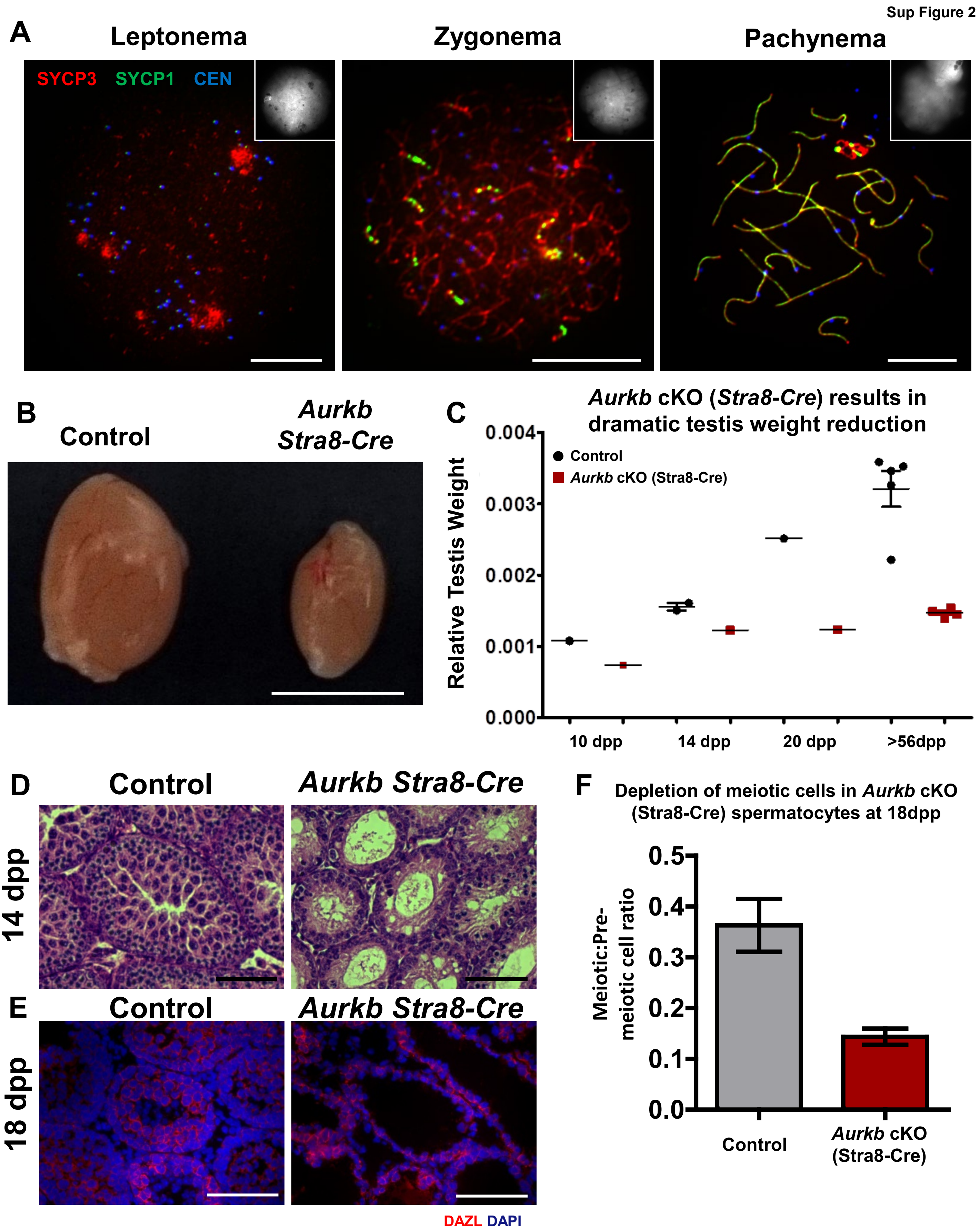
