## Supplemental Figure S3 for "Aurora B and C kinases regulate prophase exit and chromosome segregation during spermatogenesis"

A

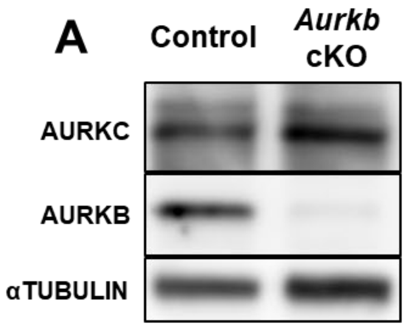


B

| Male Strain Tested | Genotype of Progeny | | Total # of Progeny | Cre Efficiency |
|---|---|---|---|---|
| | *+/Flox* | *+/Del* | | |
| *Aurkb*$^{Flox/Del}$ *Spo11-Cre* | 1 | 91 | 92 | 99% |

C D E

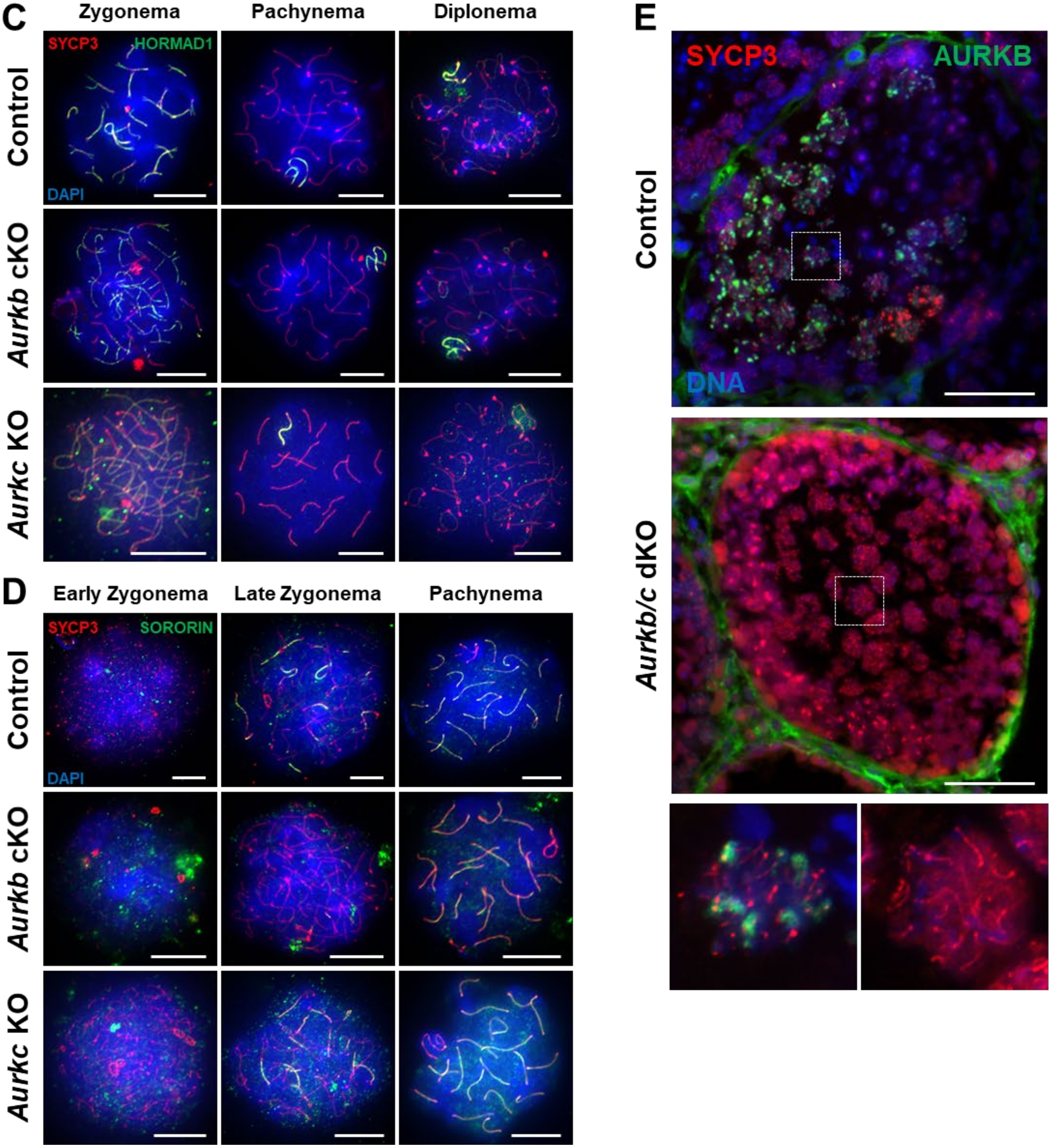
