## Supplemental Table S1 for "Aurora B and C kinases regulate prophase exit and chromosome segregation during spermatogenesis"

Sup. Table 1: Genotyping primers used in this study

| Gene | Genotyping Primers | |
|---|---|---|
| **Gene** | **Forward Primer (5'-…..-3')** | **Reverse Primer (5'-…..-3')** |
| *Aurkb* $^{tm1c}$ | AGGGCCTAATTGCCTCTTGT | GGGCATGAATTCTTGAGTCG |
| *Aurkb* $^{tm1d}$ | AGAGGTCTCCCTGCCTCTG | GGGCATGAATTCTTGAGTCG |
| *Cre Transgene* | CCATCTGCCACCAGCCAG | TCGCCATCTTCCAGCAGG |
| *Stra8-Cre* | GTGCAAGCTGAACAACAGGA | AGGGACACAGCATTGGAGTC |
| Taqman -*Neo* | CTCCTGCCGAGAAAGTATCCA | GGTCGAATGGGCAGGTAG |
| Taqman -*Csk* | CTGGCCATCCGGTACAGAAT | TGCAGAAGGGAAGGTCTTGCT |
