## Supplemental Table S2 for "Aurora B and C kinases regulate prophase exit and chromosome segregation during spermatogenesis"

Sup. Table 2: Antibodies used in this study

**Primary Antibodies**

| Antibody | Host | Source | Cat. Number | IF Dilution | WB Dilution |
|---|---|---|---|---|---|
| AIM1 (Aurora B) | Mouse | BD Biosciences | 611082 | 1:200 | 1:1000 |
| alpha-Tubulin | Mouse | Sigma | T9026 | 1:1000 | 1:10000 |
| Aurora A | Mouse | Thermo | MA5-15803 | 1:200 | |
| Aurora C | Mouse | Dr. Tang K Tang | | 1:500 | 1:2500 |
| CREST (CEN) | Human | Antibodies Incorporated | 15-235 | 1:50 | |
| gamma-H2AX | Mouse | Thermo | MA1-2022 | 1:500 | |
| H1T | Guinea Pig | Dr. Mary Ann Handel | | | |
| H3p(ser10) | Mouse | Abcam | ab14955 | 1:200 | |
| H3p(ser10) | Rabbit | Millipore | 06-570 | | 1:20000 |
| Hormad1 | Rabbit | Abcam | ab155176 | 1:200 | |
| Mad2 | Rabbit | BioLegend (Covance) | PRB-452C-200 | 1:250 | |
| Meikin | Rabbit | Dr. Yoshinori Watanabe | | 1:1000 | |
| REC8 | Rabbit | Dr. Karen Schindler | | 1:1000 | |
| SGOL2 | Rabbit | Dr. José Luis Barbero | | 1:50 | |
| Sororin | Rabbit | Dr. Susannah Rankin | | 1:150 | |
| SYCP1 | Rabbit | Thermo | PA1-16763 | 1:1000 | |
| SYCP3 | Mouse | Santa Cruz | sc-74569 | 1:50 | 1:2000 |
| SYCP3 | Rabbit | Novus | NB300-231 | 1:1000 | |

**Secondary Antibodies**

| Antibody | Host | Source | Cat. Number | IF Dilution | WB Dilution |
|---|---|---|---|---|---|
| Mouse IgG (H+L), Alexa Fluor 488 | Goat | Invitrogen | A-11001 | 1:500 | |
| Mouse IgG (H+L), Alexa Fluor 568 | Goat | Invitrogen | A-11031 | 1:500 | |
| Rabbit IgG (H+L), Alexa Fluor 488 | Goat | Invitrogen | A-11008 | 1:500 | |
| Rabbit IgG (H+L), Alexa Fluor 568 | Goat | Invitrogen | A-11011 | 1:500 | |
| Human IgG (H+L), Alexa Fluor 633 | Goat | Invitrogen | A-21091 | 1:500 | |
| Mouse IgG (H+L), HRP | Rabbit | Invitrogen | 31450 | | 1:20,000 |
| Rabbit IgG (H+L), HRP | Goat | Invitrogen | 31466 | | 1:20,000 |
